# Transcriptome Response of Cannabis (*Cannabis sativa* L.) to the Pathogenic fungus *Golovinomyces ambrosiae*

**DOI:** 10.1101/2022.08.01.501243

**Authors:** Dinesh Adhikary, Aliaa El-Mezawy, Upama Khatri-Chhetri, Limin Wu, Stephen W. Smith, Jian Zhang, Jan J. Slaski, Nat N.V. Kav, Michael K. Deyholos

**Affiliations:** Department of Agricultural, Food & Nutritional Science, University of Alberta, Edmonton, AB Canada; Bio Industrial Division, InnoTech Alberta, Vegreville, AB Canada; Ten-10 Ventures Inc, Kelowna, BC Canada; College of Agriculture, Jilin Agricultural University, Changchun, China; Department of Biology, University of British Columbia, Kelowna BC Canada

**Keywords:** *Cannabis sativa*, Powdery Mildew, *Golovinomyces ambrosiae*, Transcriptome, Biotic-stress

## Abstract

Powdery mildew (PM), caused by the obligate biotrophic fungus *Golovinomyces ambrosiae*, is an economically important fungal disease of hemp - and marijuana–type cannabis. While the PM disease can be managed effectively by cultivating resistant hosts, there is no known PM-resistant genetic variant. This is the first report of transcript level responses of the hemp cultivar ‘X59’ to *G. ambrosiae*. Transcript level changes at 5-, 8-, and 11-days post-inoculation (DPI) of *C. sativa* were evaluated against uninoculated control. Our analysis revealed that 1,898 genes were significantly (*q*-value < 0.05) differentially expressed (DE) following the pathogen challenge. Among these, 910 and 988 genes were upregulated and downregulated, respectively as the infection progressed to 11 DPI. Genes related to salicylic acid (SA), (LOC115715124 and LOC115711424) and WRKY transcription factor (LOC115707546, LOC115715968, and LOC115707511) were highly upregulated. There were 45 DEGs that were homologous to PM-related genes, including chitin elicitor receptor kinase 1 (*CERK 1*), enhanced disease resistance 2, (*EDR2*), and powdery mildew resistance (*PMR*) genes. Moreover, the genes related to glycosyl hydrolases, particularly LOC115699396, LOC115708023, LOC115710105, and LOC115710100, were highly upregulated and potentially important in mediating pathogen responses. Collectively, this study has contributed to an enhanced understanding of the molecular mechanisms that are involved in cannabis and PM disease interaction and has identified several gene candidates that can be further investigated for their role in defence mechanisms.

## 1. Introduction

Hemp (*Cannabis sativa* cv. ‘X59’) powdery mildew (PM) is caused by an obligate biotrophic fungal pathogen, *Golovinomyces ambrosiae*, affecting most of the cannabis cultivars [1]. Initially, infection appears as white mycelial growth followed by sporulation of pathogens visible as epiphytic circular patches with a white fuzzy patina on adaxial leaf surfaces [1], [2]. As the disease progresses, fungal mycelia and conidia spread to all aerial vegetative and reproductive parts, including flower bracts, buds, and stems. Apart from degrading the quality of harvested flowers and leaves, PM infection leads to leaf chlorosis, inhibits photosynthetic CO2 assimilation, reduces the capacity of infected plants to form sucrose, triggers distortion and premature leaf senescence, and eventually diminishes seedling vigour [3]–[6]

Female plants that are primarily destined for medical marijuana can be affected significantly by degraded crop quality because the value is determined by organoleptic factors such as smell and appearance of the product [7]. Leaves with PM infection show fungal mycelial growth with plenty of sporulation and spores are chiefly clustered on the sticky surface of glandular trichomes [1]. Over the course of infection, necrotic lesions are visible on the infection sites. Thus, not only is the product visually repelling but there may also be unknown health risks. While PM disease is caused by a spectrum of fungal strains, *G. ambrosiae* has been the causal agent in *Cannabis* spp. and has the highest incidence in growth facilities across different parts of Canada [8]. This has prompted the prolonged and excessive use of chemical fungicides, which again is not favorable because of residual toxicity and the tendency of enhancing selection pressure on the PM population, which can promote resistance to the fungicides [9]. Thus, the fungus *G. ambrosiae* presents a significant threat to the cannabis industry.

With the advancement of next-generation sequencing technologies, cannabis genome and transcriptome work has progressed. Although genome assemblies are available for several strains of cannabis, complete indexing of abiotic and biotic stress-responsive genes is still far from completion [8]–[10]. Transcriptome assemblies have also been generated for vegetative and reproductive tissues, focusing mostly on active metabolites such as terpenes and cannabinoids [8], [11]–[15]. Recently, Gao et al [16] and Liu et al [17] investigated the transcriptome response to drought and salinity stress, respectively, in hemp-type cultivars. McKernan et al [18] developed some preliminary information on RNA expression in response to biotic stress.

However, there is still a dearth of original research generating transcriptomic information on the PM-cannabis interaction on the species. Bearing in mind the multifaceted biotrophic nature of the PM disease, and based on the available cannabis draft genome [12], we hypothesized that the quantification of transcriptional changes, at 5-, 8-, and 11 - DPI, lead to the identification of key genes and metabolic pathways that are involved in the cannabis and *G. ambrosiae* interaction, especially in the later stages of infection. In this study, hemp (low THC cannabis) cultivar, ‘X59’ (susceptible to PM), was infected with *G. ambrosiae* and the transcriptional changes at three different time points, 5-, 8-, and 11-days post-inoculation was assessed in the inoculated and control samples at each time point. The genes identified may aid in enhancing our understanding of the potential mechanisms of cannabis and PM disease interaction and in the selection of biomarkers for further validation and investigation of their biological role in mediating defence response against the disease.

## 2. Materials and method

### 2.1 Plant Material

Seven hemp accessions (Canda, CFX2, Delores, Finola, Katani, Silesia, and X59) were obtained from the InnoTech Alberta, Vegreville germplasm collection and a preliminary disease screening was carried out. Surface sterilized seeds were sown in sterile potting mix and placed in controlled growth chamber (Conviron E15) conditions (light intensity 466 µMolm^-2^s^-1^; photoperiod 16:8 h; temperature 22 °C; and humidity 72%). Light intensity and height were adjusted as the seedling height increased. When plants were 14 days old post-germination, a pure isolate of *G. ambrosiae* was inoculated on healthy leaves. All of the inoculated seedlings developed PM infection and the symptoms were visible by 8 DPI.

### 2.2 Preparation of Fungal Pathogen and Inoculation

An isolate of *G. ambrosiae* was kindly provided by Dr. Zamir Punja (Simon Fraser University, British Columbia). The obligate biotrophic fungal isolate was cultured using 25 days old hemp seedlings under controlled growth chamber conditions as indicated above. Fungal culture was initiated 14 days prior to the inoculation of experimental plants. At least 20 young seedlings were infected with PM to culture enough fungal inoculum. By the end of day 14, fungal spores were easily visible and copiously present on the surface of culture leaves confirming the viability and virulence of the isolate. When the experimental hemp seedlings were at the 5-6 leaf stage (7 days post-germination), young leaves including developing middle leaves were lightly moistened using mist from a spray bottle. Using a soft brush, conidia were collected from infected hemp plants and dusted gently over the leaf surface of the healthy leaf. Dusting was performed close to the leaf surface ensuring that each treated plant received an equal amount of fungal inoculum. Inoculated young seedlings were covered with a plastic dome for 24 hours to maintain high humidity on the leaf surface.

### 2.3 Confirmation of Fungal Infection Using Microscopy

To confirm the fungal penetration and infection of plant tissues, asymptomatic and symptomatic leaves from 8- and 11-DPI were collected and immersed in chemical fixative, FAA (3.7% Formaldehyde, 5% Acetic Acid, and 50% Alcohol). Tissue samples were fixed for 72 hours at 4°C. Tissues were then transferred to ethanol series (50% and 70%) for drying and were embedded in paraffin blocks. Thin cross-sections of the leaves were cut using a microtome (RM2125, Leica, Wetzlar). Cross-sectioned samples were stained with toluidine blue, and then washed and mounted on a glass slide with coverslip using resinous medium. Sections were then visualized under a light microscope [Leica DMRXA microscope (Meyer Instruments, Texas)], and images were taken with QI Click digital camera and processed using Q Capture Pro 7 software (Q Imaging, British Columbia).

### 2.4 RNA Isolation

Plant tissue samples were collected and prepared following the MIQE guidelines [19]. After fungal inoculation, seedlings were inspected every 24 hours. Ensuring that the sampled leaves were of the same ages and similar developmental stages, newly opened leaves from both control and inoculated groups were collected from the two independent experiments at three different time points, 5-, 8-, and 11-DPI and flash-frozen in liquid nitrogen and kept at -80 °C until RNA extraction. Six plants were collected from experiment 1 (referred as replicate ‘1’ for each time point) and six additional plants were collected from experiment 2 (referred as replicate ‘2’ for each experiment) (Table 1). Frozen samples were homogenized using Geno/Grinder 2010 (ATS Scientific Inc. Burlington, Ontario) and RNA was isolated using RNeasy Plant Mini Kit (Qiagen, Valencia, California). To remove trace DNA, RNA samples were treated with DNAse I (Ambion, ThermoFisher Scientific, Markham, Ontario) following the manufacturer’s protocol. A total of 12 samples were prepared for RNA sequencing.

**Table 1.** RNA-Seq statistics

| Sample* | Repli-<br>cate | Read<br>Length<br>(bp) | Clean<br>Reads | Tophat<br>Mapped<br>Reads | %<br>Mapped |
| --- | --- | --- | --- | --- | --- |
| 5 DPI Control | 1 | 100 | 50,442,788 | 40,747,771 | 81 |
| 5 DPI Control | 2 | 100 | 50,238,782 | 40,132,936 | 80 |
| 5 DPI Treated | 1 | 100 | 50,330,184 | 40,611,199 | 81 |
| 5 DPI Treated | 2 | 100 | 50,408,094 | 40,869,038 | 81 |
| 8 DPI Control | 1 | 100 | 50,467,426 | 40,412,470 | 80.1 |
| 8 DPI Control | 2 | 100 | 42,009,774 | 34,733,598 | 83 |
| 8 DPI Treated | 1 | 100 | 50,664,596 | 39,380,153 | 78 |
| 8 DPI Treated | 2 | 100 | 51,100,826 | 39,079,529 | 76.5 |
| 11 DPI Control | 1 | 100 | 49,874,138 | 41,146,063 | 82.5 |
| 11 DPI Control | 2 | 100 | 50,359,286 | 41,835,600 | 83.1 |
| 11 DPI Treated | 1 | 100 | 49,030,140 | 38,728,524 | 79 |
| 11 DPI Treated | 2 | 100 | 50,620,354 | 38,637,169 | 76.4 |
| **Total** |  |  | **595,546,388** | **476,314,050** |  |
| **Average** |  |  | **49,628,866** | **39,692,837.5** | **80.13** |

\*DPI (Days Post Inoculation)

### 2.5 RNA Sequencing and Sequence Analysis

Total RNA (5-18 µg) was sent to Beijing Genome Institute (BGI Inc., Shenzhen, China) for library preparation and transcriptome sequencing. Poly(A) mRNA was isolated using oligo(dT) beads, and mRNA was fragmented into short fragments of 200 bp, which were used to synthesize the first-strand cDNA. Using a buffer mix containing dNTPs, DNA polymerase I, and RNAseH, second-strand cDNA was synthesized and purified using a QiaQuick PCR kit (Qiagen Inc., Duesseldorf, Germany). The short fragments were ligated to sequencing adapters and purified through agarose gel electrophoresis. Finally, suitable cDNA fragments from different tissue libraries were PCR amplified and sequenced using 90 bp PE Illumina HiSeq 2000 (Illumina Inc., San Diego CA, USA). Raw sequence reads were filtered to remove low quality reads and adaptor contamination and deposited in the NCBI sequence read archive (SRA) under project number PRJNA634569.

Paired end reads from all 12 libraries (Table 1) were analyzed using Tuxedo Pipeline [20]. Bowtie2 [21] was used to prepare the reference genome index and TopHat (v2.1.0) [22] to align the raw reads to the reference [12]. Mapped reads were used as the input data for reference guided transcriptome assembly and quantified differential expression using Cufflinks (v2.2.1) [20]. The GTF files generated from all 12 libraries were merged using cuffmerge. Cuffdiff was performed to assess the differentially expressed genes between the control and the treated samples at all three time points. Gene expression was assessed using Fragment Per Kilobase of transcript Per Million fragments mapped (FPKM) values and the significant differential expression for multiple comparison was assessed using Benjamini-Hochberg correction (*q* < 0.05) [23].

To validate the differential gene expression of the transcriptome data, nine putative disease resistance related genes showing a continuous pattern of either upregulation or downregulation at all three-time points were selected for qRT-PCR. Two technical replicates were used, cannabis *actin* (*CsActin)* gene that was published earlier [24] was used as the internal control to normalize the gene expression.

### 2.6 Functional Classification and Annotation of Differentially Expressed Genes (DEG)

Predicted genes were annotated using the Basic Local Algorithm Search Tool (BLASTx) to align genes to the TAIR9 protein database; two databases from Uniprot (TrEMBL and Swissprot); and National Center for Biotechnology Information (NCBI) non-redundant (nr) protein databases [25] NCBI; The UniProt Consortium, 2021]. BLAST hit results with significant homology (*e-*value, e<10^-10^, plength >150 bp, plength > 30%) were further employed for additional inferences of their biological role. TAIR9 hit IDs that correspond to the gene with significant differentially expressed values (*q*-value < 0.05, after Benjamini-Hochberg correction) were taken for gene ontology (GO) enrichment analysis using AgriGo [26]. *Arabidopsis* gene model (TAIR9) was used on the background and the following parameters were applied to run the analysis – statistical test method (Hypergeometric), multi-test adjustment statistical method [Yekutieli (FDR under dependency)], significance level (0.05), minimum number of mapping entries (5), and gene ontology type (complete GO).

## 3. Results and discussion

### 3.1 Differential Response of Hemp Cultivars to G. ambrosiae Infection

As a preliminary study, seven different hemp cultivars (Canda, CFX2, Delores, Finola, Katani, Silesia, and X59) were selected in consultation with a hemp grower (Dr. Jan Slaski, InnoTech Alberta, personal communication). All seven cultivars including ‘X59’ were challenged with *G. ambrosiae* and screened for their susceptibility to the fungus. Three parameters were carefully applied for the assessment: i) time of emergence of symptoms post-inoculation, ii) the number of leaves infected at 11 DPI, and iii) visible chlorotic lesions on the leaf surface. In the preliminary study, all tested cultivars showed fungal symptoms by 8 DPI, the number of infected leaves was variable between 2-4 in all cultivars, and the symptomatic leaves gradually turned chloritic after 12 DPI (Table S1). Thus, based on the extent of infection, all tested accessions were similar in terms of disease susceptibility to the pathogen indicating that any of the accessions would be suitable for the downstream molecular analysis. Among all others, X59 was selected as a candidate because X59 is a commercially used Canadian dual-purpose (grain-fibre type) cultivar accounting for approximately 26% of national acres in 2019 (Government of Canada). The variety was developed in early 2000 by crossing a male line (in 29) from Voronezh region and female line (in 50) from Udmurt Republic of the Russian Federation (Government of Canada).

### 3.2 PM Fungal Infection in C. sativa cv. ‘X59’

Information on the life cycle of *G. ambrosiae* is limited. Under favourable conditions, fungal conidia are produced asexually, and it takes 8-10 h for the germination [2]. Upon germination, hyphae penetrate the plant cell within 10-17 h. The whole infection process occurs within 120 h from the time a spore lands on the surface of a leaf to the point of establishment on the tissues, however, the visible infection was not observed at 5 DPI on the tested genotype (Figure 1 A1 and A2), thus the first time point was selected at 5 DPI. In this study, *G. ambrosiae* infection showed a characteristic epiphytic growth of circular patches with white fuzzy patina by 8 DPI and progressed rapidly to 11 DPI, the growth was predominantly on the adaxial surface (Figure 1 B1, B2, and C1, C2). Once the epiphytic growth was visible, the radius of the infection increased, and the onset of chlorosis on the leaf surface was observed; this symptom was likely associated with the degeneration of palisade parenchymal cells (Figure 2 A1, A2, A3, B1, and B2). As the infection progressed, infected spots were visible on the abaxial surface with an abundant colony of hyphal structures on the surface. At the microscopic level, appressorium and haustorium were visible, along with a network of hyphal structures and hyphal bridges between the cells (Figure 2 A1 and A3). Initiation of chlorosis of the leaf may have resulted due to collapsing of mesophyll palisade and spongy cells forming lumen or swollen like structures between the cells [Figure 2 A1, A3, and B1; [27], [28]]. An earlier report suggested that during PM disease development, accumulation of H_2_O_2_ first occurred in mesophyll cells that are underlying just below the infected epidermal cells and gradually H_2_O_2_ accumulated around the infected cells [28], thereby the damage to the cells might have been caused by H_2_O_2_ accumulation [29].

**Figure 1.**
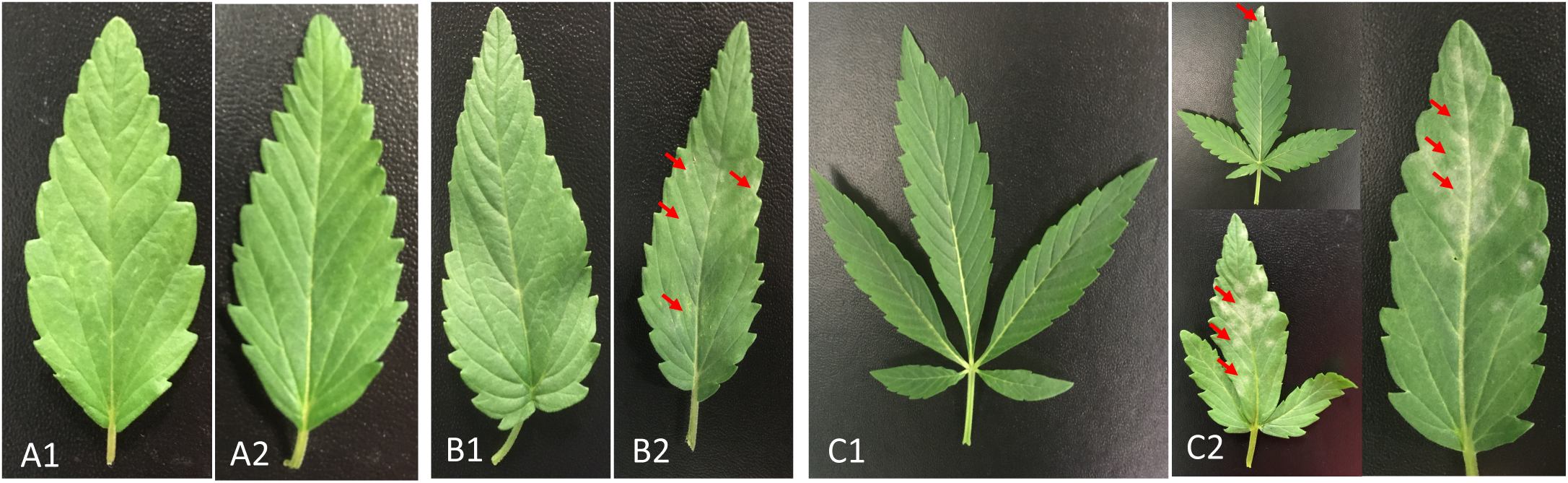
Hemp cv X59 leaf post infection. A1 and A2) Leaf sections from 5 days post inoculation (DPI). A1 and A2 represents uninoculated and inoculated leaf, respectively. B1 and B2) Leaf sections from 8 DPI. B1 and B2 represents uninoculated and inoculated leaf, respectively. C1 and C2) Leaf sections from 11 DPI. C1 and C2 represents uninoculated and inoculated leaf, respectively. Solid red arrows indicate the powdery mildew infections on the leaf surfaces.

**Figure 2.**
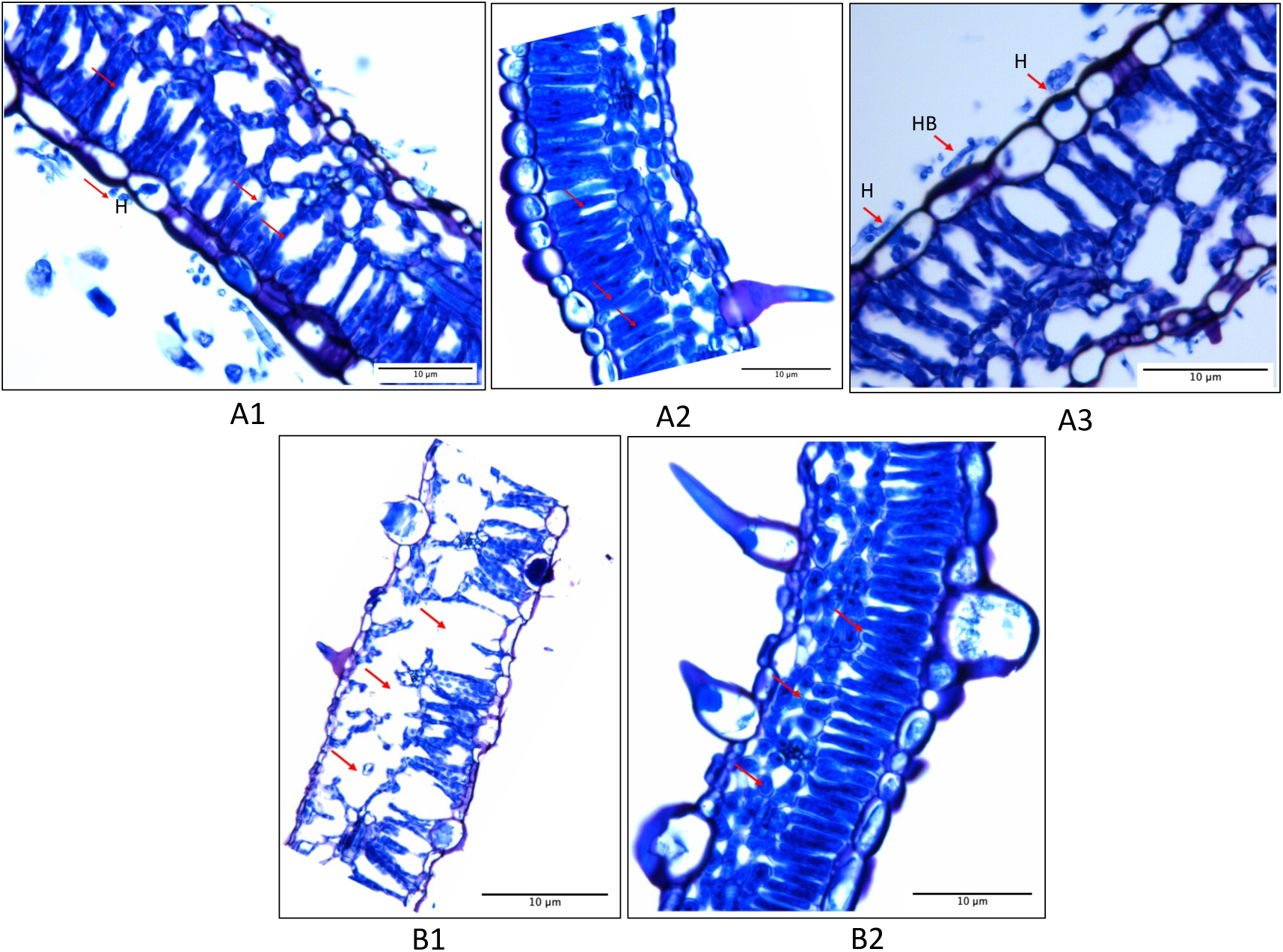
Toluidine blue stained leaf sections. A) Leaf sections from 8 DPI. A1 and A3 indicate infected tissues at 8-DPI, A2 indicates control or water inoculated tissues. B) Leaf sections from 11 DPI. B1 indicates inoculated leaf sections, B2 indicates control or water inoculated leaf sections. Red arrows indicate the modified palisade cells and big spaces between the cells; haustoria (H) and fungal hyphal bridge (HB) were also visible in the earlier stage of infection as indicated by red arrows in figure A1 and A3. Although the palisade cells have started deforming generating spaces between the cells, spongy cells look normal at this stage (A1 & A3). Both palisade and spongy cells were deformed by 11 DPI (B1); however, epidermal, palisade, and spongy parenchymal cells appear normal in the uninoculated control tissues (A2 & B2).

By 11 DPI, an early sign of chlorotic lesions was evident on the upper leaf surfaces and chlorosis on the lower surface took another 24-48 hours. Thus, the three-time points (5-, 8-, and 11-DPI) were selected to capture the transcriptomics changes (5 DPI: early stage with no visible symptoms on the leaf surface; 8 DPI: onset of visible infection on the leaf surface, and 11 DPI: full-fledged infection with visible circular patches and early onset of chlorosis).

### 3.3 RNA Sequencing and Transcript Regulation in Response to PM Infection

Reference genome-guided transcriptomic analysis was applied to a total of 12 samples generated from three-time points (5, 8, and 11 DPI) and produced 595,546,388 clean reads (Table 1). A total of 476,314,050 reads were mapped to the reference genome [12]. There were 22,762 genes that were expressed at all three-time points (i.e., with at least ten reads aligned to the reference genes). In total, 1,898 genes were significantly (*q*-value < 0.05) differentially expressed (DE) during fungal infection. Of these, 241, 315, and 910 were upregulated in the treated samples at 5, 8, and 11 DPI, respectively with the log2 fold change ranging between 1.0 and 10.0, while 263, 266, and 988 genes were downregulated at 5-, 8-, 11-DPI, respectively with a log2 fold change ranging between -1.0 and -10.0 (Table 2). After correction for multiple testing, there were 504, 681, and 1,898 significantly differentially expressed genes at 5, 8, and 11 DPI respectively with the highest number of DEGs detected at 11 DPI (Table 2) and the pattern was consistent with an earlier study in other species [30]. Several DEGs were related to the perception, recognition, and transduction of pathogen-related signals, activation of the phytohormone signaling pathway, and triggering of pathogenesis-related genes. Most of the DEGs showed consistent patterns i.e., either increasing or decreasing in the expression values at all three-time points, and the genes were identified as strong candidates for further validation and investigation of their biological role in plant defense mechanisms against powdery mildew (Table S2).

**Table 2.**
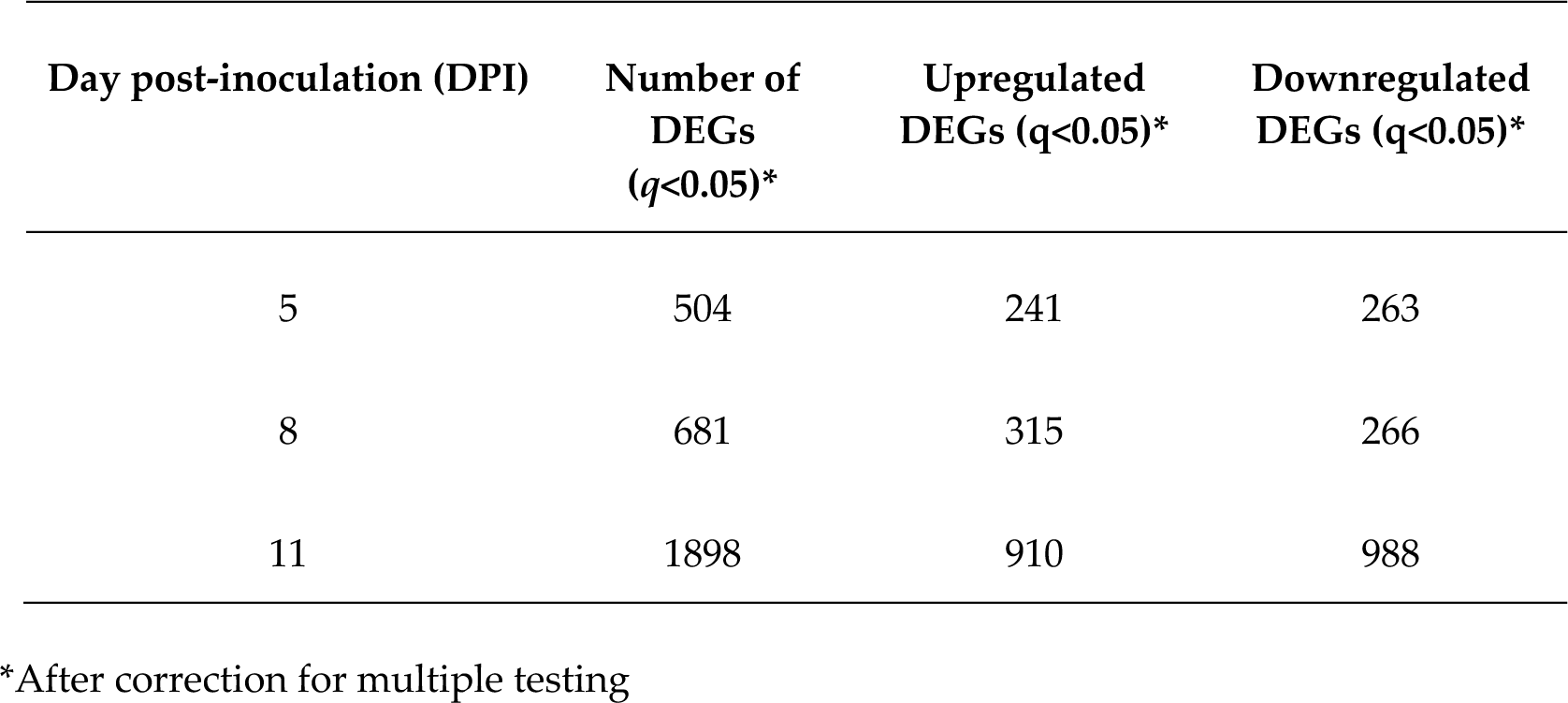
Differentially expressed genes (DEGs) detected at each time point post-inoculation

The functional study of some of these candidates is currently underway.

A few putative genes that were homologs of disease resistance-related genes, including disease resistance protein *At4g27190*-like (LOC115695607), ABC transporter G family member (LOC115714985), probable LRR receptor-like serine/threonine-protein kinase (LOC115709196), and serine/threonine-protein kinase SAPK3-like (LOC115716582) were downregulated at 5 DPI and upregulated (>3.8 log2 fold) in the later stage (11-DPI) of infection. This pattern is somewhat consistent with the gene regulation in wheat challenged with *Zymoseptoria tritici* where many genes related to disease resistance were downregulated in the earlier infection stages and upregulated in the later stages [31]. Both *G. ambrosiae* and *Z. tritici* are biotrophic in nature and it is possible that when the infection becomes more severe, their signal transduction system is fine-tuned, and it is also likely that the host is capitalizing more resources on the synthesis of defense-related gene and proteins, thereby the expression level changes as the infection progressed to later stages.

RNA-seq results were validated by performing qRT-PCR on nine genes that were differentially expressed (Figure 3; Table S3). Gene candidates for qRT-PCR were selected randomly, and explicitly for the validation of the two techniques. All the tested genes had similar patterns of gene expression between RNA-seq and qRT-PCR log2 fold change data (Figure 3).

**Figure 3.**
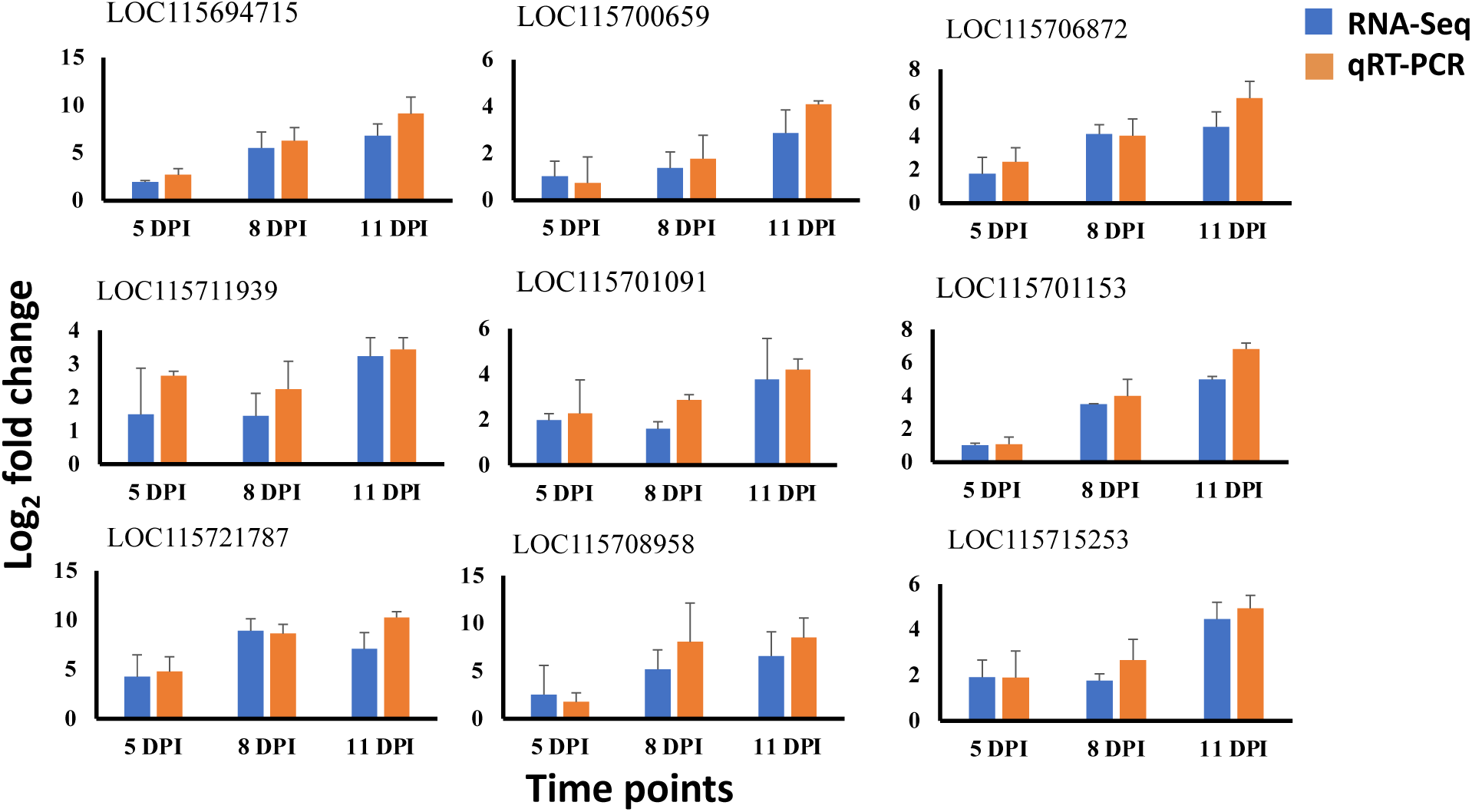
Comparison of gene expression pattern, qRT-PCR vs RNAseq. From the pool of RNAseq expression values, nine putative disease resistance related genes were selected and used for qRT-PCR validation. The ids starting with ‘LOC11…’ represent the gene ids corresponding to the original IDs in the reference genome [12]. Their corresponding gene sequences can be obtained from- https://ftp.ncbi.nlm.nih.gov/genomes/all/annotation_releases/3483/100/GCF_900626175.1_cs10/. Y-axis represents the log2 fold change and the x-axis represents the time points after the fungal inoculation. Error bar represents the standard error mean.

### 3.4 Gene Ontology Enrichment Analysis

By the end of 11 DPI, 81 unique GO terms were significantly (FDR <0.05) enriched (Table S4); some of the highest-level categories included cellular (GO:0009987), metabolic process (GO:0044237), and biosynthetic process (GO:0009058), response to – stimulus (GO:0050896), – stress (GO:0006950), and - biotic stimulus (GO:0009607). There were categories directly related to host-pathogen interactions such as response to -fungus (GO:0009620), - chitin (GO:0010200), - bacterium (GO:0042742), and - external stimulus (GO:0009605). Associated with these categories, there were several DEGs annotated as chitinases, resistance genes containing TNL (TIR-NBS-LRR) domains, and powdery mildew specific genes (Figure 4; Table S4). Categories such as cellular- (GO:0044237), primary- (GO:0044238), secondary-metabolic process (GO:0019748), and phenylpropanoid biosynthetic pathways (GO:0009699) were some of the highest-level enriched categories in metabolic processes. Other highly enriched categories included protein metabolic process (GO:0019538), biosynthetic processes (GO:0009058), and response to hormone stimulus (GO:0009725) (Table S4). The enrichment of hormone-related categories reflects the fundamental role of phytohormones in the plant-pathogen interaction and defense system. Phytohormones such as salicylic acid (SA), jasmonic acid (JA), and ethylene (ET) play a crucial role in regulating physiological processes, including plant immunity and the primary defense against fungal pathogens [32].

**Figure 4.**
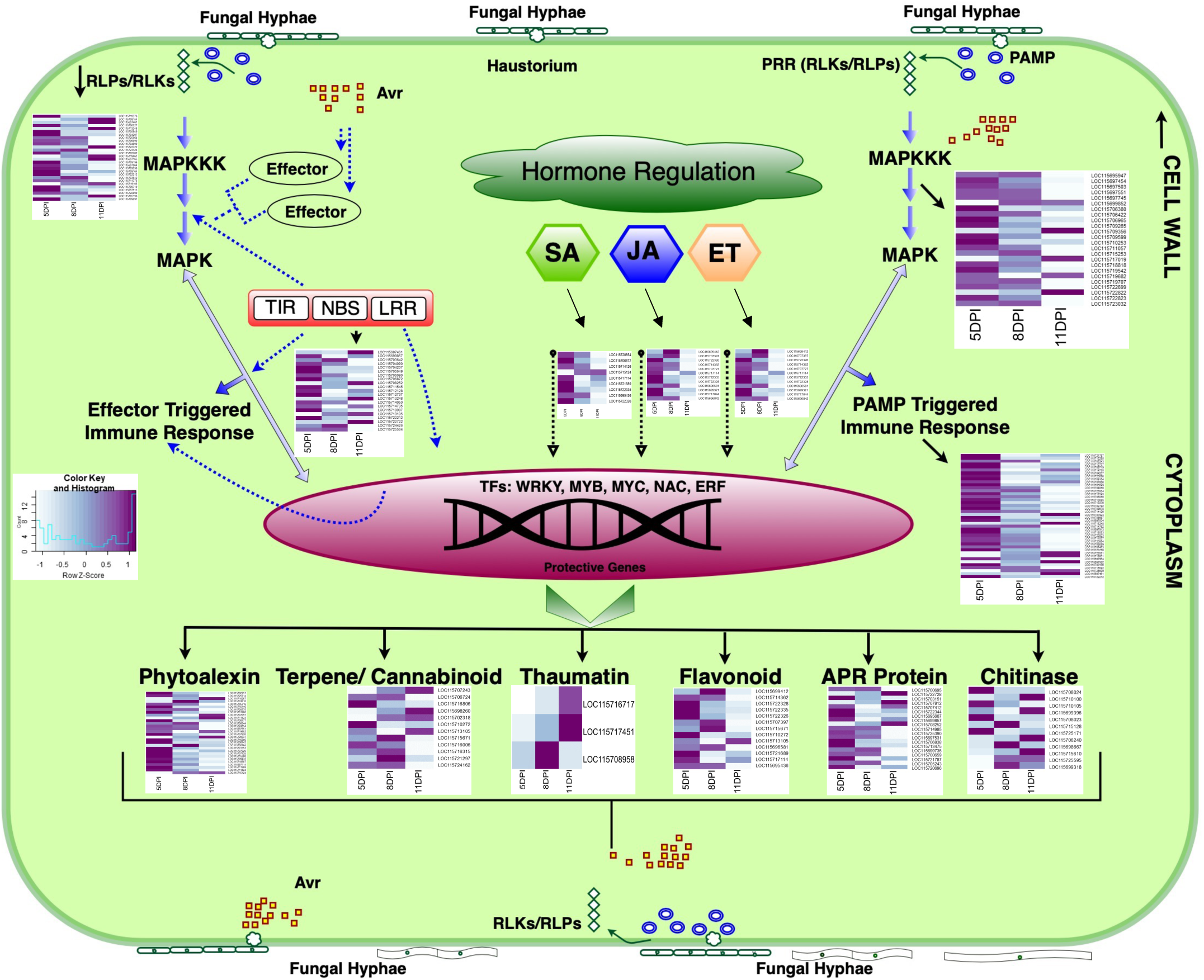
Schematic diagram showing plant-fungal interaction upon PM inoculation in cannabis. Heatmaps for log2-fold change values at 5-, 8-, and 11-DPI were shown besides each major molecular steps in the plant-pathogen interaction pathway. Dark purple indicates downregulation and light color indicates upregulation. Putative genes that were shown in the heatmap were significantly differentially expressed at least at one time point (*q*<0.05). During the course of fungal infection, pathogen produces elicitors such as PAMPs and effectors, upon perceiving the foreign molecular patterns, host receptors RLKs/ RLPs and R genes interact with the elicitors, and the interaction leads to the activation of primary signal transduction pathway such as MAP kinase. Eventually, transcription factors are regulated and a whole host of defense related molecular changes takes place, including activation of phytohormone (SA, JA, and ET) biosynthetic pathway and secondary metabolism related genes. Glycosyl hydrolases (chitinase and β-1, 3-glucanase) acts on the degradation of fungal cell wall; likewise, thaumatin and defensin permeabilize membrane bilayers of microbial cells and interfere with the microbial protein synthesis, other low molecular mass organic compounds such as phytoalexin inhibit the establishment and metabolism of pathogen activities, thereby inhibiting fungal establishment, colonization, and infection on the host cells. Moreover, secondary metabolites such as flavonoids and terpenes play a vital role in the antioxidant activity against reactive oxygen species produced during pathogen invasion. Avr = Avirulence factor; RLP = Receptor like proteins; RLK = Receptor like kinase; SA = Salicylic acid; JA = Jasmonic acid; ET = Ethylene; PRR = Pathogen recognition receptor; PAMP = Pathogen associated molecular pattern; MAPK = mitogen-activated protein kinase; TIR = toll/interleukin-1 receptor; NBS = nucleotide binding site; LRR = leucine rich repeat APR = adult plant resistance

The GO functional categories of external stimulus (GO:0009605) and response to other organisms (GO:0051707) included some plant defence-related genes. Among the upregulated genes were several genes homologous to receptors of pathogen signals such as CNL (CC-NBS-LRR), RLK (receptor-like kinases), RLP (receptor-like proteins), and TNL (TIR-NBS-LRR) (Figure 4). Specific genes such as chitin elicitor receptor kinase 1 *(CERK1),* enhanced disease resistance 2, *(EDR2)*, and powdery mildew resistant 5 *(PMR5)* protein) were highly upregulated. Many of these genes and several others are discussed in more detail below in reference to Figure 4.

### 3.5 Resistance Genes (R-genes)

To date, over 200 R genes in *Arabidopsis* have been reported (http://www.prgdb.org/prgdb/plants/?id=151). Seven types of domains have been defined within them: nucleotide-binding site (NBS), leucine-rich repeat (LRR), toll/interleukin-1 receptor (TIR), coiled-coil (CC), serine-threonine kinase (STK), transmembrane (TM), and leucine zipper (LZ). These domains have been combined into seven categories namely, CC (coiled-coil), CNL (CC-NBS-LRR), RLK (receptor-like kinases), RLP (receptor-like proteins), Pto (Ser/Thr kinase protein), NLR (NBS-LRR), and TNL (TIR-NBS-LRR) [33]–[37]. In this study, there were 11 putative TNL genes, 11 RLP*s*, and 4 RLK-like genes that were differentially expressed (Table S5-A). Twenty-three of them were upregulated and three of them were downregulated by 11 DPI. Among the receptor-like proteins, six putative genes (LOC115700527, LOC115704090, LOC115704207, LOC115705649, LOC115706090, and LOC115725564) that were homologs of receptor-like protein 52 (RLP52) were consistently upregulated by 11 DPI. Given that R genes recognize pathogen effectors and play a vital role in gene-for-gene interaction, they are identified as crucial members of the plant immune system. These surveillance gene transcripts are expected to be expressed consistently even at low levels in response to the pathogen; however, two of the genes, LOC115713248 (receptor-like protein kinase 2) and LOC115705057 (protein too many mouths) were downregulated as the infection progressed to 11 DPI. The decrease in transcript abundance of R genes is modulated by the host micro-RNA and it is observed in the absence of a pathogen interference [38]. However, we saw quite the opposite for the two genes. Although it is unclear why the genes were downregulated, it is likely indicating the susceptibility factor, which is a subject of further exploration. Similar observations were made in flax transcriptome study, flax seedlings infected with fungal pathogen also reported the downregulation of the two similar R genes [39]. Furthermore, a predicted gene (LOC115708252) homologous to disease resistance RPP13-like protein, containing a CNL domain, was consistently upregulated >3.0 log2 fold by 11 DPI, indicating that the gene is strongly involved in the host-pathogen interaction. An earlier report has demonstrated that *RPP13* provides resistance against downy mildew [40]. Similarly, RPP13-like protein was reported to play a vital role in the perception of bacterial effectors; its loss of function led to susceptibility to biotrophic microbe (*Xanthomonas campestris* pv. *campestris*) and diminished resistance to *P. syringae* [35], [41]–[43]. Thus, the gene (LOC115708252) is potentially important for further exploration of the resistance mechanism in cannabis. Similarly, among the TNL-related genes, four of them (LOC115711545, LOC115712128, LOC115712737, and LOC115706872) were expressed >4.0 log2 fold. Taken together, this study revealed several putative genes homologous to *RLP52, RPP13*, and other CNL / TNL domains that are potentially involved in the plant-pathogen response and are suitable candidates for further investigation of the resistance mechanism. However, not all of the R genes can contribute equally to durable resistance against PM, thus combinations of multiple R genes and other genes such as adult plant resistance (APR) genes (discussed below) into a single plant have been demonstrated effective in gaining durable resistance against fungal diseases in cereal crops [44]. There is a potential that some of the identified R and APR genes, upon validation of their biological role, particularly those that were upregulated or downregulated consistently at all three-time points, can be combined using the gene pyramiding method through traditional breeding practices or modern genomics tools such as genome editing and develop elite cannabis variety that can confer durable resistance to the pathogen.

### 3.6 Mitogen-Activated Protein Kinase (MAPK) Signaling in Response to PM Infection

The mitogen-activated protein kinase (MAPK) cascade is a primary signal transduction pathway in higher eukaryotes and plays a vital role in plant development and pathogen response [45]–[48]. Plants employ multi-step defense responses against pathogens, beginning with activation of the kinase upon perceiving PAMPs/MAMPs and pathogen effectors [49]. In the present study, 13 kinases were differentially expressed representing *MAPKKK1, MAPKKK5*, *MAPK3*, *MAPK6*, *MAPKKK NPK1*, and serine/threonine-protein kinase *EDR1* (Figure 4; Table S5 - B) and eleven of them were upregulated. A predicted gene (LOC115712285), homologous to *MAPKKK* serine/threonine-protein kinase *EDR1* was increased > 5.0 log2 fold. Earlier studies have demonstrated that *EDR1* plays a key role in the regulation of MAP cascade and aids in the sensing and conferring resistance against fungal diseases including powdery mildew [48]–[52]. Similarly, two predicted genes (LOC115711057 and LOC115719682), homologous to mitogen-activated protein kinase 3 (*MPK3)*, were consistently upregulated as the infection progressed to 11 DPI (Figure 4). An earlier study on *MPK3* demonstrated that upon *B. cinerea* infection, *MPK3* was activated and induced camalexin synthesis in *Arabidopsis* [53]. Likewise, Asai et al [34] demonstrated that the activation of the MAPK cascade provided resistance against both fungus and bacterial pathogens. Thus, it is reasonable to speculate that the upregulated genes in the pathogen-challenged groups are strongly associated with cannabis and PM disease interaction, and these genes are potential candidates for further functional characterization and investigation of biological role in mediating resistance response against the PM disease.

### 3.7 Transcription factor (TFs) in Response to PM Infection

In the natural environment, plants are unremittingly exposed to abiotic and biotic stressors, which affect plant growth and development. Upon pathogen invasion, transcriptional reprogramming occurs, and transcription factors (TFs) are employed in the defense signaling mechanism, interacting, and binding with stress-related genes in a sequence-specific manner orchestrating the gene expression in response to the microbial invader [54]. In this study, 243 plant TFs were differentially expressed, 172 of them were upregulated and 71 were downregulated (Table S6). Among the TFs, six major TF families, *bHLH*, *ERF, MYB, NAC, WRKY*, and *bZIP* were well represented in the data set (Table S6) and these TFs were demonstrated highly regulated in response to biotic and abiotic stress response [54]–[56]. As the master regulators of plant defense response, these TFs were either up or downregulated in different plant species.

For instance, the *bZIP* TF showed prominent activation in response to *Ustilago maydis* infection in maize [57]. In the current study, there were a total of eight *bZI*P TFs that were differentially expressed; of them, four genes were upregulated, and the remaining four were consistently downregulated at 5-, 8-, and 11-DPI (Table S6). Similarly, the WRKY protein family with the signature conserved domain, WRKY-DBD, plays a vital role in the recognition of the W-Box element and positively modulating and activating the early defense-related genes such as PAMP responsive genes [57]–[60]. There was a total of 23 WRKY-related genes that were differentially expressed. Of them, 18 were upregulated, while four were downregulated. Six genes were expressed >5 log2 fold due to the pathogen interaction (Table S6). Some of the differentially expressed putative genes, LOC115707546 (homologous to *WRKY46*), LOC115715968 (homologous to *WRKY53*), and LOC115707511 (homologous to WRKY DNA-binding transcription factor 70) were linked to the positive regulation of resistance against *Pseudomonas syringae* in *Arabidopsi*s [61]. While the transcriptional reprogramming of *WRKY53* precisely regulates oxidative responses to both biotic and abiotic stresses, *WRKY46* and *WRKY70* together enhance resistance to *P. syringae* potentially by increasing the expression of salicylic acid (SA) dependent genes [61] [62]. Given that *WRKY53* plays a fundamental role in response to both biotic and abiotic stresses in wheat and rice, overexpression of the TF has enhanced the accumulation of PR proteins and reduced the infection from rice blast fungus, *Magnaporthe oryzae* [63]. Notably, the two genes (LOC115715968 and LOC115707511) homologous to *WRKY53* and *WRKY70*, increased expression >7 log2 fold in response to the *G. ambrosiae* (Table S6) and the transcript abundance of the genes increased gradually in the treated groups with the progression of infection, indicating that the genes were strongly involved in the PM and cannabis interaction. Thus, all three genes (LOC115707546, LOC115715968, and LOC115707511) are considered as potential candidates for further investigation against the PM disease in cannabis.

Similar to the WRKY transcription factor families, some MYB TFs form a complex network of regulatory responses against the biotic stress [64]. In response to *G. ambrosiae,* 32 genes were differentially expressed, seven genes were downregulated, and 25 were upregulated; of which, five putative genes (LOC115721787, LOC115705224, LOC115705243, LOC115711509, and LOC115711509) have transcript abundance >5 log2 fold (Table S6). Earlier studies showed that *MYB44* was involved in defense response against *Alternaria brassicicola* in *Arabidopsis*, *AtMYB44* regulated transcriptional activation of *WRKY70* and induced defense response against *A. brassicicola* [65]. The presumed homolog of *MYB44* (LOC115711127) showed transcript abundance gradually increased >3 log2 fold as the disease progressed to 11 DPI (Table S6).

While most of the TFs discussed above were upregulated, there were seven genes related to MYB that were downregulated consistently at all three-time points in response to the pathogen indicating a potential susceptibility factor (Table S6). MYB related TFs, for instance, *myb46* knockdown mutant showed increased resistance against *B. cinerea* [66]. Moreover, there were other TFs with genes that were downregulated, *WRKY* (four genes), *NAC* domain (three genes), *bHLH* (two genes), *bZIP* (four genes), and *ERF* (two genes) (Table S6). Although TFs constitute a large family of proteins and the functional role of many of them is yet to be characterized, some of the earlier studies on TFs have demonstrated both the positive and negative roles in the plant immunity response [67]–[69]. For instance, *NAC* is one of the well-studied TFs and they were confirmed as both positive and negative regulators of defense-related genes [70]. Many of the TFs are induced under pathogen influence and play a vital role in linking signal transduction processes between defense-related phytohormones and ROS-related pathways during plant and pathogen interaction. Overexpressing *NAC6* and *NAC111* in rice have enhanced tolerance to rice blast and bacterial blight, respectively [71], [72]. Similarly, *NAC122* and *NAC131* were induced by *Magnaporthe griesa* infection and demonstrated a positive regulatory response against the rice blast resistance [73]. On the contrary, overexpression of *NAC4* has led to increased cell death and damage of cell membrane in rice [70]. Similarly, *NAC069* demonstrated a negative regulatory role in lettuce; however, when silenced, enhanced the resistance trait against *Pseudomonas cichorii* bacteria [74]. Taken together, several of the identified genes were strongly involved in cannabis interaction and PM disease interaction and they will be strong candidates for further investigation of defense mechanism; while some of the DEGs that were activated during interaction could also reflect induced disease susceptibility factor, which is again a subject of further functional validation.

### 3.8 Genes Related to Secondary Metabolic Pathways in Response to PM Infection

Secondary metabolites serve a range of vital functions in plants including defense roles [75]. Some flavonoids and terpenoids have been reported to exhibit antimicrobial properties [76]. Terpenoids from oregano oil showed promising antifungal activity [77]. Similarly, several cannabinoids have shown potent antimicrobial activity [78]. Thus, these secondary metabolites play a crucial role in the defense role against microbial pathogens including biotrophic fungus. In the current study, 13 genes related to flavonoid synthesis were upregulated, 10 of them upregulated >3.0 fold by 11 DPI (Figure 4; Table S5-C). Five homologs of flavanone 3-dioxygenase (LOC115714362, LOC115722328, LOC115722326, LOC115707397, and LOC115696581) were consistently upregulated as the infection progressed to 11 DPI. Transcript, LOC115722326, was upregulated >5 log2 fold. The substrate(s) of the enzyme encoded by this flavanone 3-hydroxylase (F3H) genes is unknown; however, a previous study has shown that the F3H converts *2S*-naringenin to (*2R, 3R)-* dihydrokaempferol [79]. Moreover, in another study, the compound was highly expressed in response to the infection caused by *Endoconidiophora polonica* [80]. The enzyme is also in the biosynthetic pathway for catechin, another pathogen-defensive molecule [80]. Furthermore, among the enriched GO terms, the secondary metabolic process (GO:0019748) was one of the highly enriched terms potentially indicating the importance of genes that were involved in the biosynthesis of metabolites in response to the PM infection.

There were eight terpene-related genes that were significantly differentially expressed following *G. ambrosiae* infection (Table S5-D). Homologs of terpene synthase, geranylgeranyl pyrophosphate synthase, farnesyl pyrophosphate synthase 1, and germacrene B synthases were well-represented in the dataset and most of them were upregulated by 11 DPI. Notably, one of the putative terpene synthase genes, LOC115716806, was consistently upregulated at all three time points and showed > 8.0 log2 fold by 11 DPI. Earlier studies have showed that overexpression of terpene synthase in rice conferred enhanced resistance to the fungus *Magnaporthe oryzae* [81].

Some cannabinoids have shown potent antimicrobial activity [78]. Our data showed that nine genes putatively associated with the cannabinoid biosynthesis pathway were significantly differentially expressed (Figure 4; Table S5-E). Of these, two were downregulated and seven were upregulated. Although there is a dearth of scientific evidence to support the antifungal properties of cannabinoids *in planta*, limited *in vitro* studies have shown that some of the active cannabinoids have shown antifungal activities against *Cryptococcus neoforms* and *Candida albicans* [80]–[84].

Thus, the differential expression of the putative genes involved in the secondary metabolic pathways indicates the plant response to the pathogen. Many of the cannabis homologs especially the upregulated putative genes are potentially linked to the plant and pathogen interaction and can be suitable biomarkers to further investigate the defense mechanism against the pathogen.

### 3.9 Phytoalexin Synthesis and Regulation In Response to PM Infection

Phytoalexins are low molecular mass organic compounds that are produced against pathogens and inhibit their establishment, metabolism, and development in a host plant [85]. In the present study, 36 genes that were related to phytoalexin were differentially expressed, 32 of them were consistently upregulated at all three-time points with 8 putative genes that were >5.0 log2 fold change. Two genes (LOC115707097 and LOC115708777) that were homologous to indole acetaldoxime dehydratase (*CYP71A13*) were upregulated consistently at all three-time points (Table S5-F). Plants carrying the *cyp71A13* mutation were highly repressed in the production of camalexin upon challenged by *Pseudomonas syringae*; however, when the mutants were supplied with indole-3-acetonitrile exogenously, camalexin synthesis was restored. Thus, it was concluded that the homolog was potentially involved in conferring resistance against the pathogen [86]. Similarly, two genes (LOC115725714 and LOC115702757) homologous to dolabradiene monooxygenase were continually upregulated > 2.0 log2 fold by 11 DPI. Earlier study in maize demonstrated that the transcript accumulation for the homolog was increased in response to two fungal pathogens (*Fusarium verticillioides* and *F. gramenearum*) [45]. Finally, two putative genes (LOC115698743 and LOC115698987) homologous to momilactone synthase was differentially expressed at all time points. The transcript abundance in the control samples was very low to undetectable at all three-time points, while the transcript gradually increased in the treated samples (Figure 4; Table S5-F). An earlier study in rice demonstrated that the homolog was involved in the chemical synthesis of momilactone phytoalexins [47] and this diterpenoid secondary metabolite plays an essential role in the plant-pathogen interaction. Thus, the putative genes that showed consistent expression patterns at all three-time points in response to the pathogen indicate a strong biomarker for further investigation of cannabis and PM interaction.

### 3.10 Hormone Regulation in Response to PM Infection

Soon after pathogen perception, transcriptional changes trigger hormone signaling [87]. There are multiple phytohormones involved in the defense response, primarily salicylic acid (SA), jasmonic acid (JA), and ethylene (ET) [88] are the key players. Our study revealed 13 SA-related genes that were differentially expressed, and all 13 DEGs were upregulated consistently at all three-time points. There were five genes (LOC115706872, LOC115721689, LOC115722335, LOC115695436, and LOC115722326) that were upregulated >3 log2 fold by 11 DPI (Figure 4; Table S5-G) and the transcript abundance increased as the infection progressed to 11 DPI (Figure 4; Table S5-G). Of them, one putative gene, LOC115720854 [homologous to calmodulin-binding protein 60 G (*CBP60G*)] was linked to the transcription activation of SA pathway genes and other defense related genes in *Arabidopsis;* notably, overexpression of *CBP60G* in *Arabidopsis* contributed to the SA accumulation, microbe-associated molecular patterns (MAMPs) recognition and subsequently enhanced resistance to *P*. *syringae* [89]. There are several genes associated with the accumulation of SA, genes such as phytoalexin deficient 4 *(PAD4),* enhanced disease susceptibility 1 *(EDS1)* were involved in the SA regulation [90]. The predicted genes, LOC115715124 (homologous to *PAD4-like*), and LOC115711424 (homologous to *EDS1-like*) were upregulated >3 log2 fold change by 11 DPI (Figure 4; Table S5-F). Moreover, homologs of other positive SA regulators, such as LOC115715663 and LOC115700633 [homologous to nonrace-specific disease resistance 1 *(NDR1)]* and LOC115706872 and LOC115725742 [homologous to suppressor of *npr1-1* constitutive 1 *(SNC1)],* were differentially expressed and the transcript abundance for these putative genes increased by 11 DPI (Figure 4; Table S5-F). There were five DEGs representing *DLO1* genes (LOC115717114, LOC115721689, LOC115722335, LOC115695436, and LOC115722326), where the latter four were upregulated > 4 folds, although they were not involved in the pathogen resistance and were regarded as partially redundant and suppressor of immunity, these genes were differentially expressed indicating strong host response to the fungus and could be a potential susceptible factor in response to the pathogen (Table S5-G) [91]. Additionally, GO enrichment showed phenylpropanoid biosynthetic process (GO:0009699), phenylpropanoid metabolic process (GO:0009698), and response to hormone stimulus (GO:0009725) as the most enriched categories (Table S4). Given that SA pathway is well studied and there are several transcriptomic studies where SA related biosynthetic processes were enriched and genes were upregulated in response to fungal diseases including powdery mildew [30], [92]. Information revealed from this study aligns well with those earlier findings indicating that the genes identified can be strong candidate for further exploration of *G. ambrosiae* and cannabis interaction in mediating resistance against the pathogen.

Along with SA, both ET and JA play a crucial role in the activation of defense system against pathogens. Assessment of ET biosynthetic pathway revealed 15 ET-related genes that were upregulated by the 11 DPI with four putative genes (LOC115722326, LOC115707727, LOC115696581, and LOC115695436) with expression level >4 log2 fold change (Figure 4; Table S5-H). Two genes (LOC115717044 and LOC115696842) represented the 1-aminocyclopropane-1-carboxylic acid (*ACC*) synthase (*ACS*), a rate-limiting enzyme of ET biosynthesis pathway, and 11 genes represented 1-aminocyclopropane-1-carboxylate oxidase *(ACO)* and two genes represented ethylene-responsive transcription factor 1B *(ERF1B)* and are upregulated by >4 log2 fold change at the 11 DPI (Figure 4; Table S5-H). The activation of these ET-related genes by plant host upon fungal challenge was reported in an earlier study in flax [39]. While ET plays a vital role in the plant developmental process and in plant-biotic response, a comprehensive role of ET synthesis during biotic interactions is poorly understood, ET can also function as a negative signaling factor during host and pathogen interaction. Zhao et al [93] demonstrated that *rice dwarf virus-induced* ethylene production by stimulating S-adenosyl-L-methionine synthetase, a key player in ET biosynthesis, in rice. Thus, upregulation of ET-related genes is an indication of successful pathogen interaction, but it may not always indicate a resistance factor. However, *ERF1B*, a transcriptional activator, is one of the crucial genetic markers involved in plant defense response against necrotizing fungus (*Botrytis cinerea* and *Plectosphaerella cucumerina*) [94], [95], and earlier studies have revealed that the overexpression of the transcription factor conferred resistance against broad range of necrotizing and soil borne fungal pathogens [39], [91]. Although all of the ethylene related genes detected in the study were upregulated in response to the pathogen, it is of note that *ERFs* and other ET related genes are also versatile in nature and they are induced not only in response to biotic stress, but also their upregulation implies developmental changes and orchestration of progress in pathogen in the tissue level [96].

JA is also involved in the defense against plant pathogens [97]. Our study revealed 20 putative JA-related genes that were differentially expressed, and a majority of these were upregulated by 11 DPI (Table S5-I). Given that JA is a well-studied defense hormone and genes that were primarily involved in the JA synthesis, allene oxide cyclase (*AOC)*, lipoxygenase 2 (*LOX2)*, and allene oxide synthase (*AOS*) were identified and their abundance was gradually increased over the course of time and upregulated >2 fold by 11 DPI (Table S5-I). Moreover, there were ten genes homologous to cytochrome P45094B3 (*CYP94B3*) and the majority of them were upregulated by 11 DPI. Earlier studies have shown that *CYP94B3* employs negative feedback control mediating catabolism of jasmonyl-L-isoleucine [98]. Concurrent transcript upregulation of both JA synthesis genes and their repressors were also found in *Arabidopsis* [55]. This indicates the balance of JA production potentially maintains the excess levels of the compound in the plant system [56], [97], [98]. Taken together, the genes identified in this study demonstrate clear alignment with earlier studies, which validates that the data is reliable; and it is safe to speculate that the genes identified in this study can potentially be a dependable source to further investigate cannabis and PM disease interaction, which is still far from a thorough investigation.

### 3.11 Glycosyl Hydrolases in Response to PM Infection

Plants respond to fungal pathogens by producing enzymes such as chitinase and β-1, 3-glucanase, which dissolve components of a fungal cell wall, such as chitin and β-1, 3-glucan [99]. The degradation of fungal cellular components inhibits microbial establishment and colonization on a plant host; moreover, pathogen-associated molecular patterns (PAMPs) are readily available to plant pattern recognition receptors (PRRs), thus preventing the microbes from entering the host. In this study, there were 14 differentially expressed genes that were the homologs of chitinase and glucanase genes (Figure 4; Table S5-J). There were three homologs of endochitinase EP3-like genes (LOC115708024, LOC115699396, and LOC115708023) that were putatively involved in the endo-hydrolysis of chitin molecules [100]. Transcript abundance of all three genes gradually increased and for the latter two, log2 fold change was > 4.50 by 11 DPI. An earlier transcriptomic study in cannabis demonstrated that under PM infection, among several other genes, chitinase-related genes were differentially expressed indicating their potential role in the pathogen response [18]. Furthermore, this study identified seven putative glucan endo-1, 3-β-glucosidase genes that were differentially expressed, five of them were downregulated, and two genes (LOC115710105 and LOC115710100) were upregulated > 5.0 log2 fold change by 11 DPI. Some of these putative glucosidase genes showed contrasting changes in transcript abundance (Figure 4; Table S5-J). In earlier studies, overexpression of glucanase gene (*ScGluD2*) in *Nicotiana benthamiana* conferred resistance against *Ralstonia solanacearum* and *Botrytis cinera* [101]. In contrast, when *Arabidopsis* seedlings were challenged with a novel strain (30C02) of cyst nematode, relative expression of glucan endo-1,3-β-glucosidase (*At4G16260*) declined after 3-5 days post root infection [102]. Further investigation on the *At4G16260* mutant showed increased susceptibility to the pathogen while overexpression of the gene conferred improved tolerance to the pathogen indicating that the gene was involved in the pathogen response [102]. Thus, the upregulated homologs of chitinases and glucanases indicate a strong interaction between *G. ambrosiae* and cannabis and potentially be suitable biomarkers for further investigation of the plant defense response against the pathogen.

### 3.12 Thaumatin in Response to PM Infection

Defensin- and thaumatin-like proteins (TLPs) interfere with microbial protein synthesis and their functions, thereby inhibiting pathogen infection to the host plant [18]. There were 31 genes with significant BLAST hits representing thaumatin related genes (data not shown), however, only three of these were significantly differentially expressed in response to *G. ambrosiae* with one of the genes (LOC115716717) that was homologous to thaumatin-like protein upregulated by >5 log2 fold change (Figure 4; Table S5-K). Similarly, two other genes (LOC115717451 and LOC115708958) representing thaumatin-like proteins were upregulated (>2 log2 fold). Similar to this study, when cannabis was challenged with PM disease, 23 TLPs were revealed indicating that TLPs play a crucial role during plant and PM interaction [18]. These antifungal thaumatin-like proteins lyse microbial cells by forming transmembrane pores in the fungal membrane [103]–[106]. Albeit several of these TLPs are yet to be functionally validated, many of the genes encoding these antimicrobial peptides are associated with defense mechanism in other plant species. Thus, homologs identified in this study are potential biomarkers to further investigate defense mechanisms in cannabis and PM interaction.

### 3.13 Adult Plant Resistance (APR) Genes in Response to PM Infection

Plants carry numerous resistance genes coding for immune receptors (TNL or CNL) and they are activated in all parts of the plant and are effective at all developmental stages; however, there are specific resistance genes that are expressed later in the adult stage towards maturity and these are categorized as the adult plant resistance (APR) genes [107]. Although APR genes represent a minority of known R genes, these genes have been applied in wheat breeding programs for decades [107]. In this study, there were 16 genes that were homologous to four different APR genes, *Lr22a, Lr34, Lr67*, and *Xa21,* and were differentially expressed. Of these, 12 genes were upregulated and 4 of them were downregulated (Figure 4; Table S5-L). Based on the protein sequence similarity, top hit putative homologs were selected and were similar to the APR gene sequence.

Although the underlying mechanism of APR genes on the prolonged resistance is yet to be understood, elite wheat varieties that were developed by stacking R genes and APR genes such as *Lr34* have provided prolonged resistance against leaf rust and powdery mildew in wheat [108]. Similarly, *Xa21* has provided resistance against leaf blight in rice [107], [108], and *Lr22a* and *Lr67* have provided resistance against leaf rust in wheat and barley [109]–[111]. *Lr67* is known to provide a multi-pathogen resistance [107], cannabis genes (LOC115707412 and LOC115722344) that are homologous to *Lr67* have strong sequence similarity to the homologs and were upregulated at all three-time points (Figure 4). Likewise, there were four genes (LOC115700659, LOC115720696, LOC115705243, and LOC115721787) that were homologous to *Lr34*; the latter two were upregulated > 5 log2 fold change by 11 DPI. Homologues of *Lr34* have also been demonstrated to provide broad-spectrum resistance against leaf rust, stripe rust (*Yr18*), stem rust (*Sr57*), and powdery mildew (*pm38)* in wheat [112]–[115]. Although all the referenced homologs were from monocots, they have strong sequence similarity thus the putative cannabis APR genes can potentially be suitable candidates for further functional validation and subsequently develop suitable biomarkers and investigate the defense mechanism in cannabis against PM causing pathogens.

### 3.14 Powdery Mildew and Bud Rot Related Genes

In the present study, there were 45 differentially expressed genes homologous to 14 unique genes that have previously been investigated for their resistance against powdery mildew and bud rot [25], [116]–[129] (Table S7). There were three putative genes (LOC115704207, LOC115719169, and LOC115725564) homologous to *AT3G21630* [chitin elicitor receptor kinase 1 (*CERK1)*], all of which were consistently upregulated >3.0 log2 fold by 11 DPI (Table S7). Earlier studies in *Arabidopsis* demonstrated that a mutation in *AT3G21630* impeded the activation of a majority of chitooligosaccharides-responsive genes and led to higher vulnerability to biotrophic fungus (*G. ambrosiae*) and necrotrophic fungus (*A. brassicicola)* [122]. Likewise, the knockout mutants of the homolog could not respond to fungal MAMPs in plants and were highly susceptible to fungus [130]. When the mutants were complemented with their wildtype (WT) gene copy, plants were able to recover WT *CERK1* function and induced the production of ROS in response to the fungal chitin elicitor [130]. Another putative gene worth noting was LOC115719467, homologous to enhanced disease resistance 2, (*EDR2*). An earlier study in *Arabidopsis* demonstrated that the homolog negatively regulated defense mechanism that was induced by powdery mildew. When *edr2* mutants were challenged by fungal pathogen, tissue chlorosis followed by necrosis was observed; however, the cell death response was localized and did not spread beyond the inoculated points indicating that the disruption phenotype was favorable to gain resistance against biotrophic fungus such as *G. ambrosiae*. Similarly, there were two genes (LOC115701063 and LOC115699540) homologous to *PMR5* (AT5G58600) in *Arabidopsis thaliana,* and both the genes were upregulated consistently by 11 DPI (Figure 5). In the earlier study, the mutant phenotype of the *pmr5* gene conferred resistance against powdery mildew caused by *G. ambrosiae* and *Erysiphe orontii* [131]. Additionally, mutants carried an enriched pectin layer, and smaller cell size indicating defects in cell expansion. Moreover, double mutants of *pmr5* and *pmr6* demonstrated a strong reduction in cell size along with high induction of uronic acid indicating that the double mutant affected pectin composition in the cell [131]. Both the mutants were important in the host cell wall modification and for effective plant-fungal interactions. While the homologs of *CERK1, EDR2,* and *PMR4* in cannabis are yet to be functionally validated, overexpression of *CERK1*, and loss-of-function mutation of *EDR2,* and *PMR4* may potentially involve in cannabis and PM interaction and the genes are strong candidates for further investigation.

**Figure 5.**
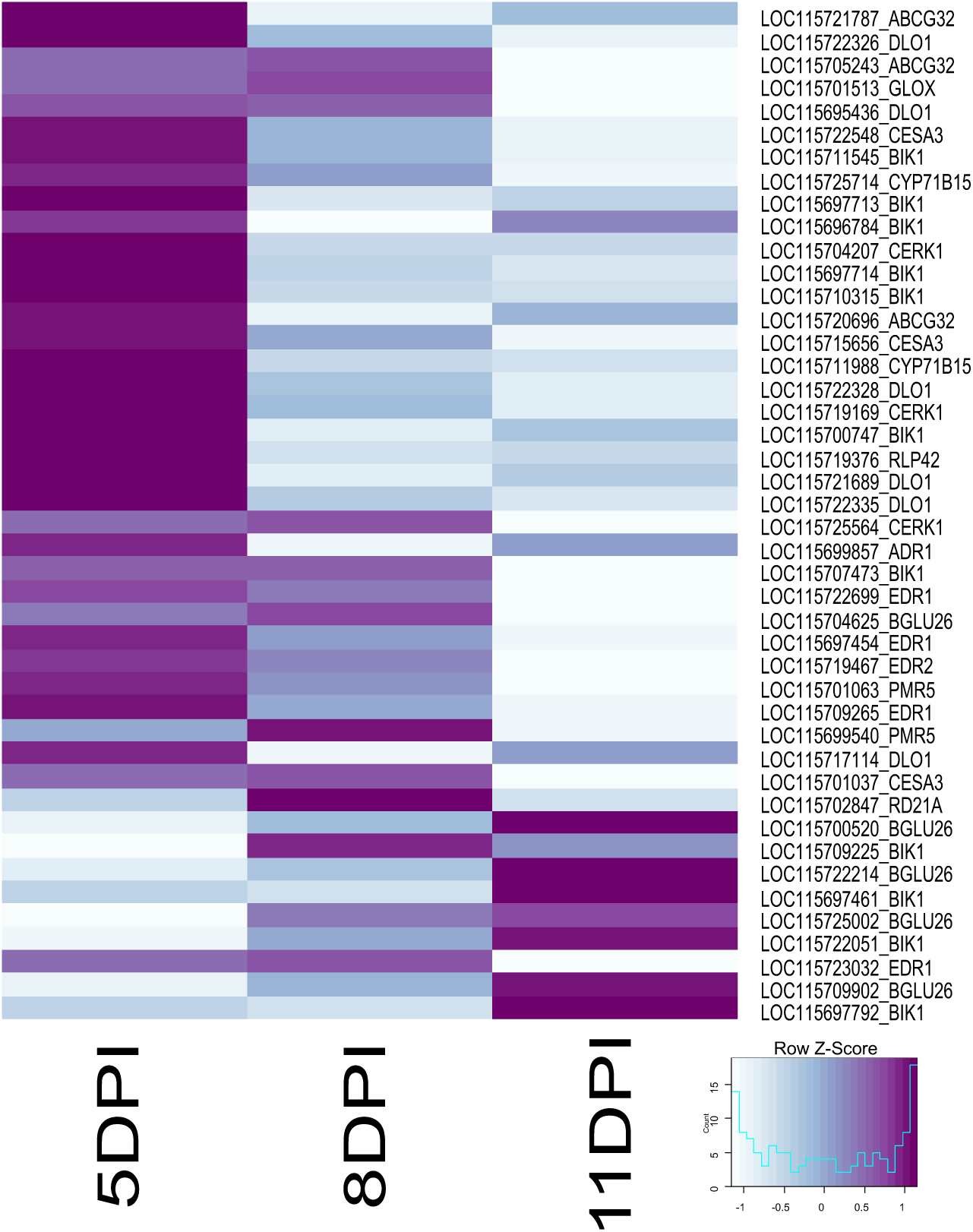
Heatmap showing the gene expression pattern of disease-specific genes in hemp through the course of time after the inoculation with *G. cichoracearum* infection. Putative genes that were shown in the heatmap were significantly differentially expressed at least at one time points (*q<*0.05). The ids starting with ‘LOC11…’ represent the gene ids corresponding to the original IDs in the reference genome [12]. Their corresponding gene sequences can be obtained from- https://ftp.ncbi.nlm.nih.gov/genomes/all/annotation_releases/3483/100/GCF_900626175.1_cs10/. Dark purple indicate downregulated and light color indicates upregulation. Full length description of the annotation can be obtained in Table S7.

### 3.15 Mildew Resistance Locus (MLO)

Mildew Resistance Locus O (MLO) related genes, particularly loss-of-function mutants, are known to confer resistance against powdery mildew in several monocot and dicot plants, such as wheat [132], tomato [133], pea [134], and arabidopsis [135]. Recently, Mckernan et al [18] reported MLO genes in cannabis. In the current study, there were 16 MLO-related genes annotated as feronia receptor-like protein kinase, and all of them were consistently upregulated at all three-time points as the infection progressed to 11 DPI (Table S8). Mutants of *feronia* (*fer*) have been previously shown to increase resistance to PM [136]. In resistant genotypes, secretory vesicles attenuate pathogen penetration by reinforcing the cell wall, and this is usually associated with MLO mediated resistance [137]. However, normal MLO genes function as negative regulators of the secretory vesicle-associated defence system, thereby making a host susceptible to the PM fungus [138]. In the current study, many MLO related genes, including LOC115719491, LOC115702328, LOC115712737, and LOC115709196 showed log2 fold change > 3.0 in response to the pathogen and can be potential candidates for further functional validation using genome editing and investigate their potential role in mediating defence response against PM fungus.

## 4. Conclusions

To our knowledge, this is the first transcriptomic-wide report on hemp cultivar ‘X59’ - *G. ambrosiae* interaction. In recent years, the cannabis industry is rapidly growing, and breeders are shifting towards marker-assisted breeding, single nucleotide polymorphisms (SNP), and quantitative trait loci (QTL) mapping for improving agronomic properties for a multitude of applications including disease resistance. As a preliminary step towards elucidation of the molecular basis for host and PM disease interaction, we have employed RNAseq and developed comprehensive transcriptome information. These results have contributed to a better understanding of the transcriptional changes involved in cannabis responses to the PM causative fungus and led to the identification of several key genes and metabolic pathways that are potentially involved in the host and the fungus interaction. It is supported by the upregulation of genes related to thaumatin, glycosyl hydrolases, phytoalexin, flavonoids, and phytohormones. SA-related genes particularly *PAD4,* and *EDS1* seem important in triggering host response to the pathogen, and glycosyl hydrolases particularly chitinases and glucanases involved in endo-hydrolysis of chitin molecules seem vital in responding against the pathogen. At the time of this investigation, neither PM-resistant cannabis genotypes nor real and near-isogenic lines for the cultivar of interest were available, precluding a comparison of gene expression in resistant versus susceptible plants. Despite this, our study has identified several genes that are important during PM disease development, including 45 genes that were potentially involved in fungal resistance against bud root and PM-related diseases and these genes will potentially be strong candidates for further validation of their biological role in mediating resistance response against the PM disease in cannabis. This study has opened avenues for further exploration of specific *Avr* genes in response to the pathogen, and other plant defence-related genes involved in the overall plant immunity. Additionally, some of the putative genes especially with high upregulation or downregulation expression values are still unannotated, thus they require further evaluation and characterization using overexpression and knockout mutants and elucidate their biological role. The transcriptome information developed in this study will also be a valuable resource for annotating the cannabis genome, which is still developing. Nevertheless, there is still a need for omics information on the host-pathogen interaction developed from real isogenic or near-isogenic lines, which would potentially make a better comparison, especially when targeting specific pathogen response genes.

## Notes

### Competing Interest Statement

The authors have declared no competing interest.

