## Supplementary material for "Transcriptome Response of Cannabis (*Cannabis sativa* L.) to the Pathogenic fungus *Golovinomyces ambrosiae*": Table S1

**Table S1.** Infection status of powdery mildew (PM) in the tested varieties**.
*** DPI = Days post inoculation

| **Hemp Variety** | **CFX-2** | **Delaros** | **Finola** | **Katani** | **Silesia** | **X59** | **Canda** |
| --- | --- | --- | --- | --- | --- | --- | --- |
| PM Infection | Yes | Yes | Yes | Yes | Yes | Yes | Yes |
| Emergence of Infection at | 5 DPI | 5 DPI | 5 DPI | 5 DPI | 5 DPI | 7-8 DPI | 5-7 DPI |
| Number of Infected Leaf | 3-5 | 4 | 5 | 3 | 3 | 2-4 | 4 |
| Visible Chlorotic lesions | 12 DPI | 12 DPI | 12 DPI | 12 DPI | 12 DPI | 12 DPI | 12 DPI |
| Disease Susceptibility | Susceptible | Susceptible | Susceptible | Susceptible | Susceptible | Susceptible | Susceptible |
